## Supporting Information for "4’-Fluorouridine mitigates lethal infection with pandemic human and highly pathogenic avian influenza viruses"

### Supporting information captions

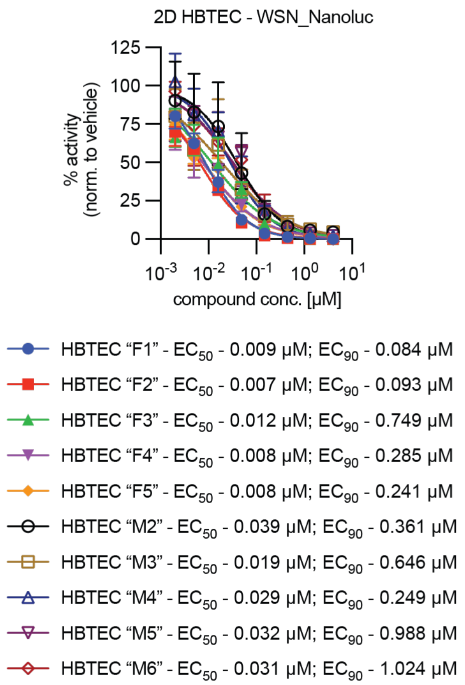


**S1 Figure:** 4’-FlU antiviral activity in 2D HBTEC. Dose-response assays of 4’-FlU against A/WSN/33 (H1N1) on HBTEC derived from 10 different healthy donors, five male and five female. Lines represent 4-parameter variable slope regression models; symbols represent data means ± SD; n=6. $EC_{50}$ concentrations and 95% CI are specified.

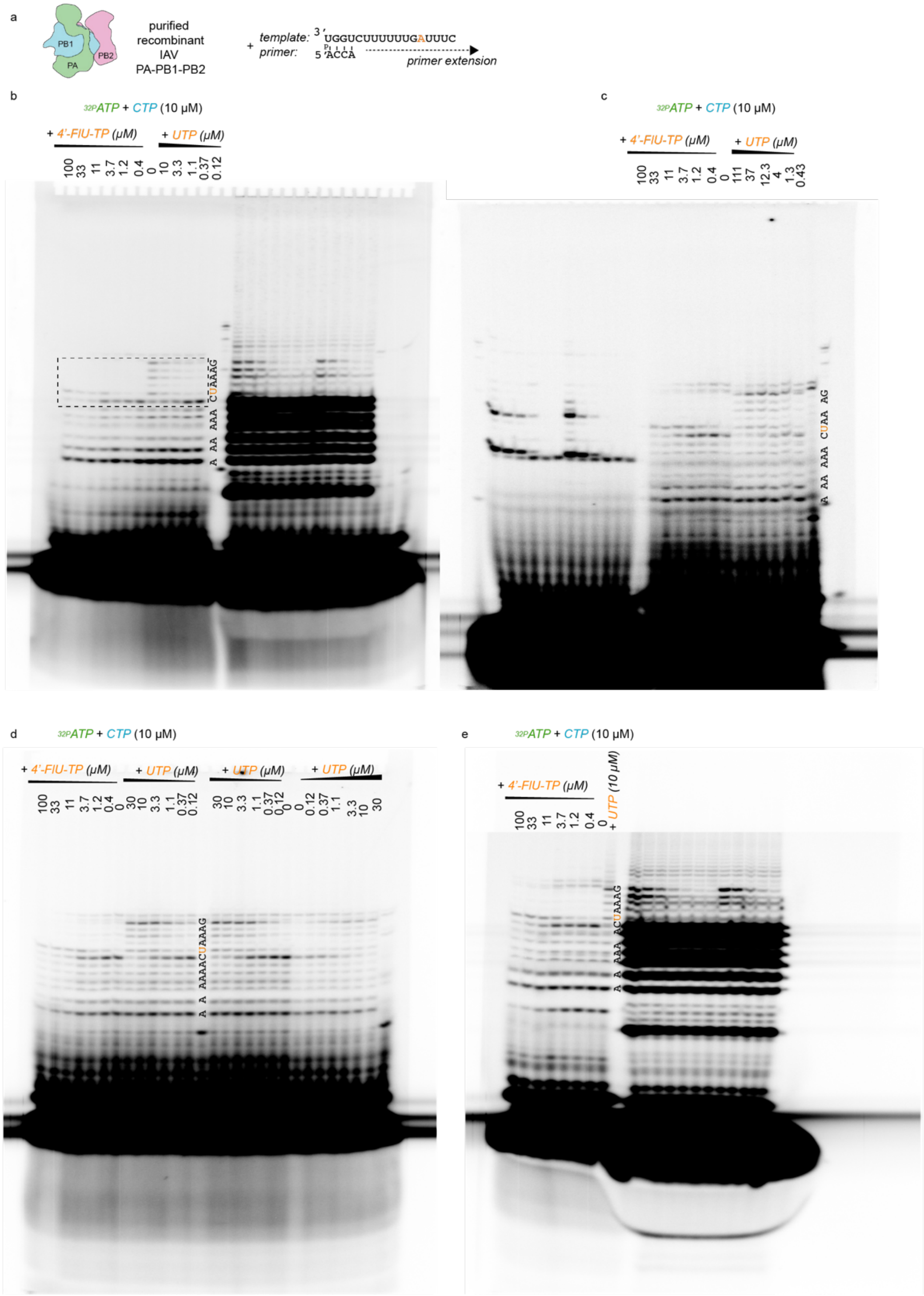


**S2 Figure:** Urea-PAGE fractionation (uncropped) of RNA transcripts. **a)** Template and primer used in the reaction. **b-e)** Independent repeats of primer extension assay by IAV polymerase. Dashed rectangle shows insert presented in figure 1e. Replicates were included in quantitations presented in figure 1e-f.

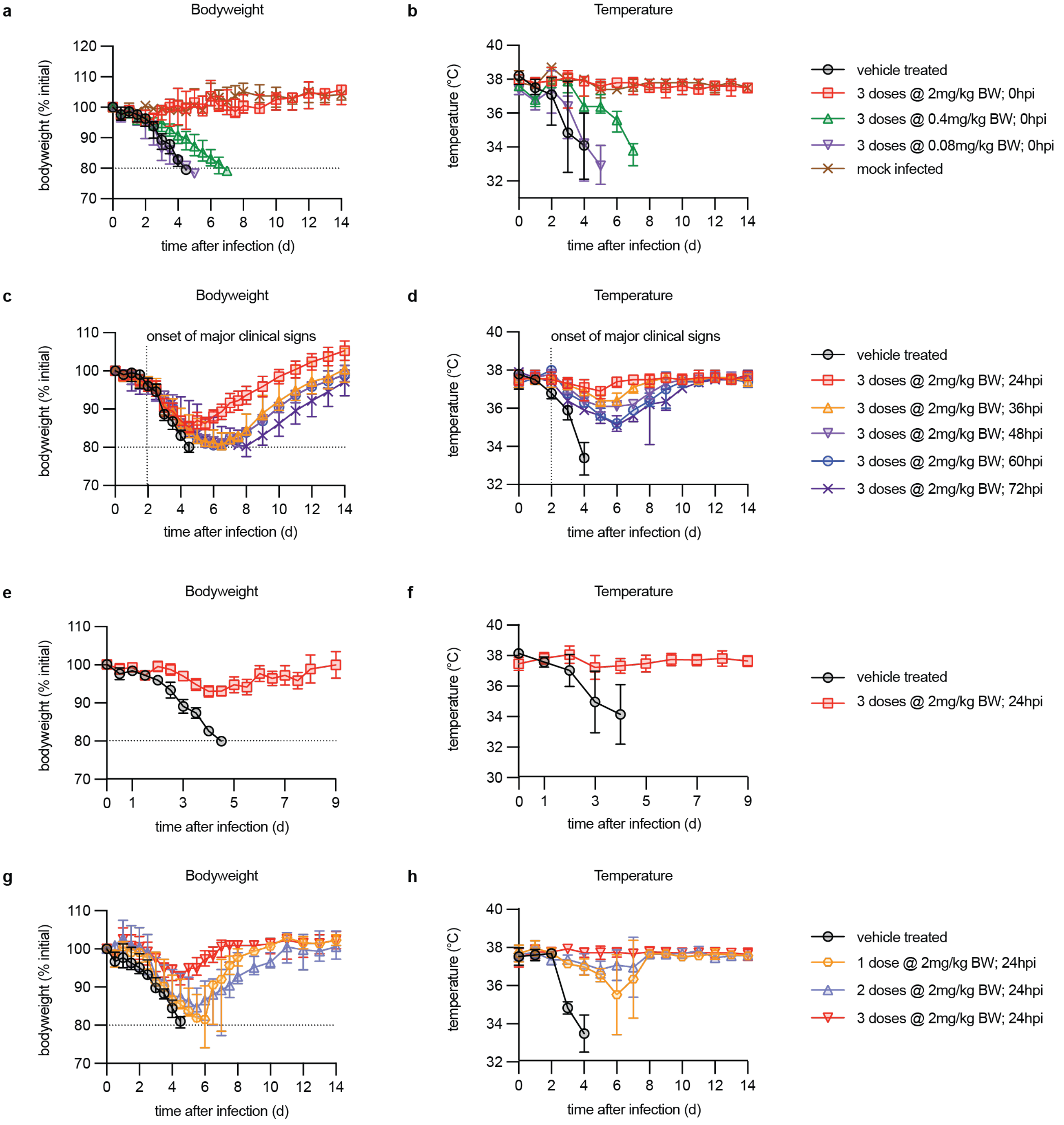


**S3 Figure**: Clinical signs in 4’-FIU-treated mice used to establish treatment paradigms. **a-h)** Body weight measurements taken twice daily (a, c, e, g) and rectal body temperature determined once daily (b, d, f, h). Dashed horizontal line specifies humane endpoint of 20% body weight loss; symbols represent data means ± SD; n values are specified in main figure 2. Results are shown for lowest efficacious dose (a-b), latest onset of efficacious treatment (c-d), virus clearance (e-f) and minimal number of doses required (g-h) studies.

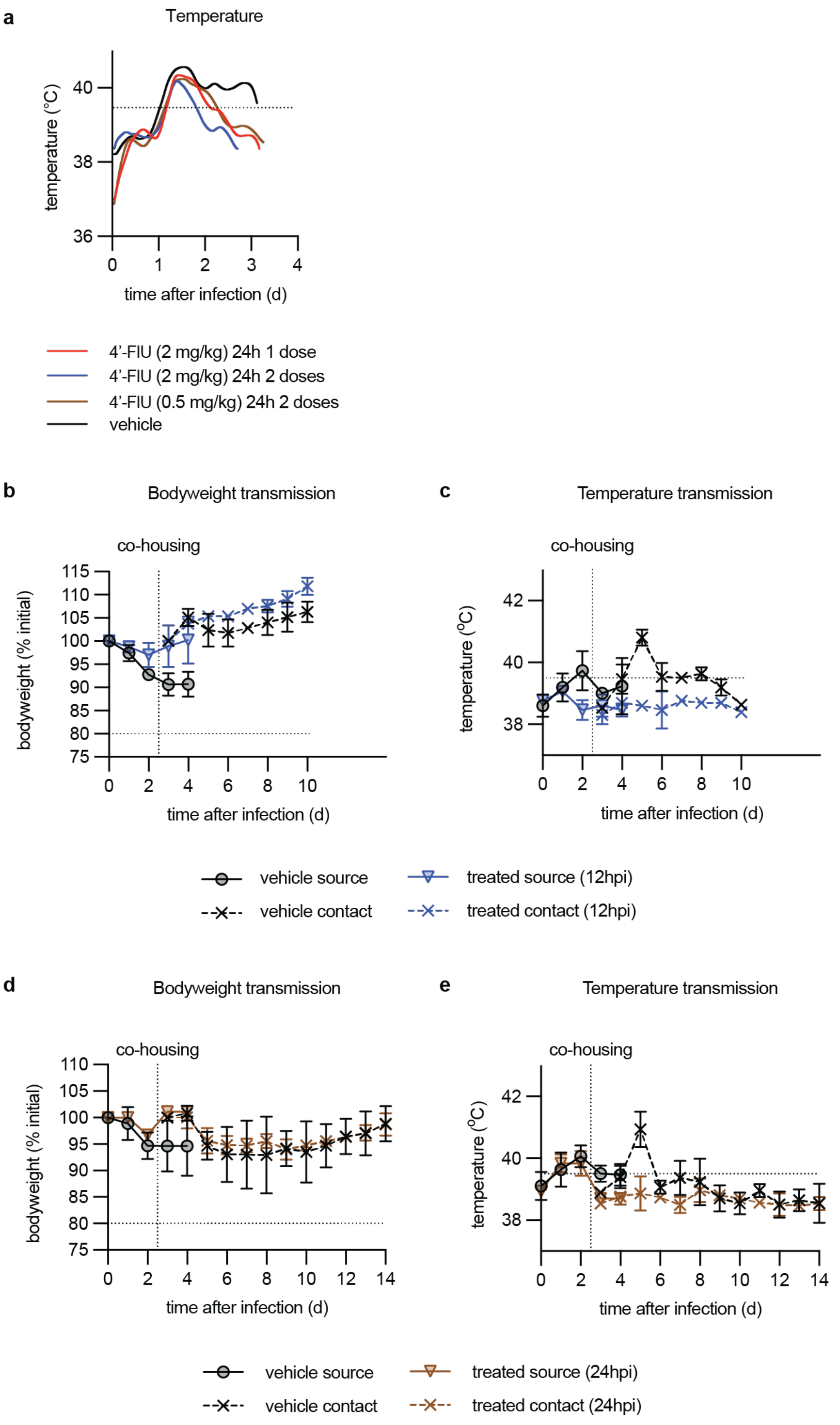


**S4 Figure:** Clinical signs in 4'-FIU treated ferrets. **a)** Continuous telemetric assessment of body temperature of ferrets in the 4'-FIU efficacy study. **b-e)** Once-daily body weight (b, d) and rectal body temperature (c, e) of ferrets involved in the Ca09 transmission studies.

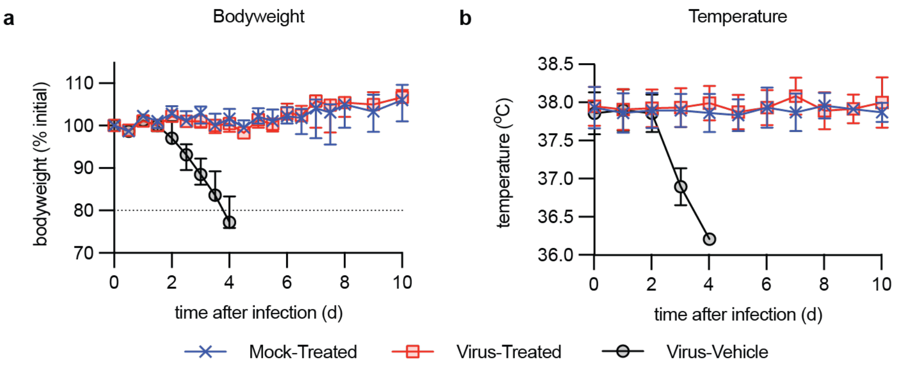


**S5 Figure:** Clinical signs in 4'-FIU-treated mice used in immune response-mitigation study. **a-b)** Body weight measurements taken twice daily (a), and rectal body temperature determined once daily (b). Symbols represent data means ± SD; n values are specified in figure 4.

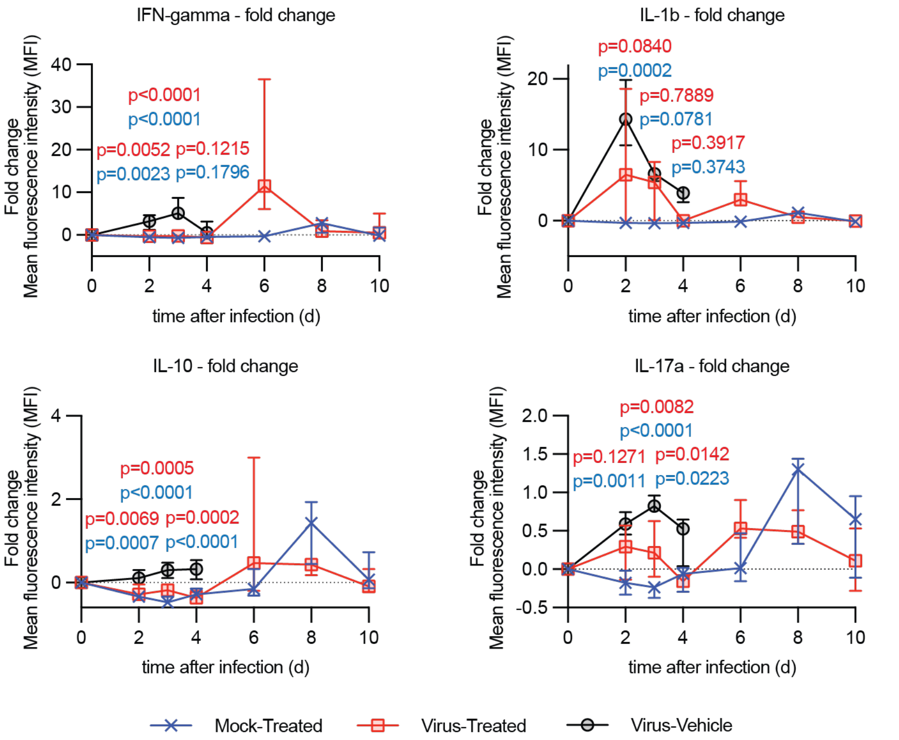


**S6 Figure:** Changes in select BALF cytokine levels in infected and 4'-FIU-treated mice. IFNy, IL-1b, IL-10, and IL-17a levels present in BALF of animals treated as in figure 4a are shown, calculated relative to levels at time of infection. Symbols represent data medians with 95% CI; 2-way ANOVA with Tukey's *post hoc* test; P values are specified.

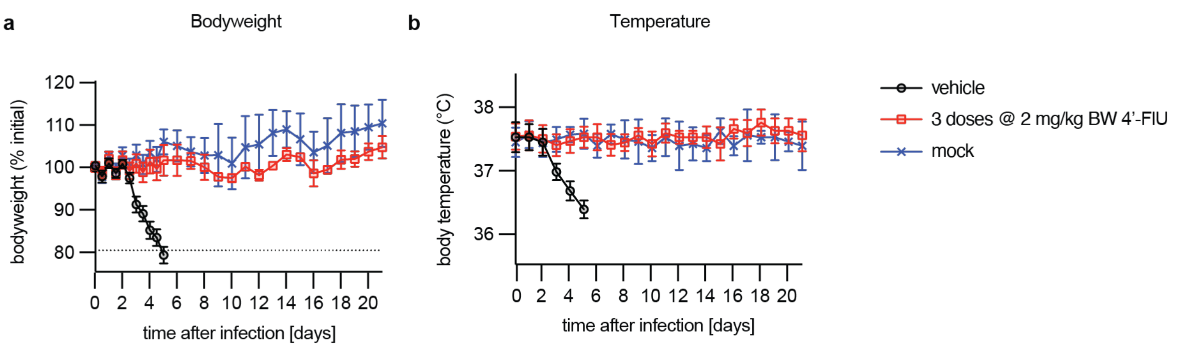


**S7 Figure:** Clinical signs of mice involved in mitigation of histopathology study. **a-b)** Body weight measurements taken twice daily (a), and rectal body temperature determined once daily (b). Symbols represent data means ± SD.

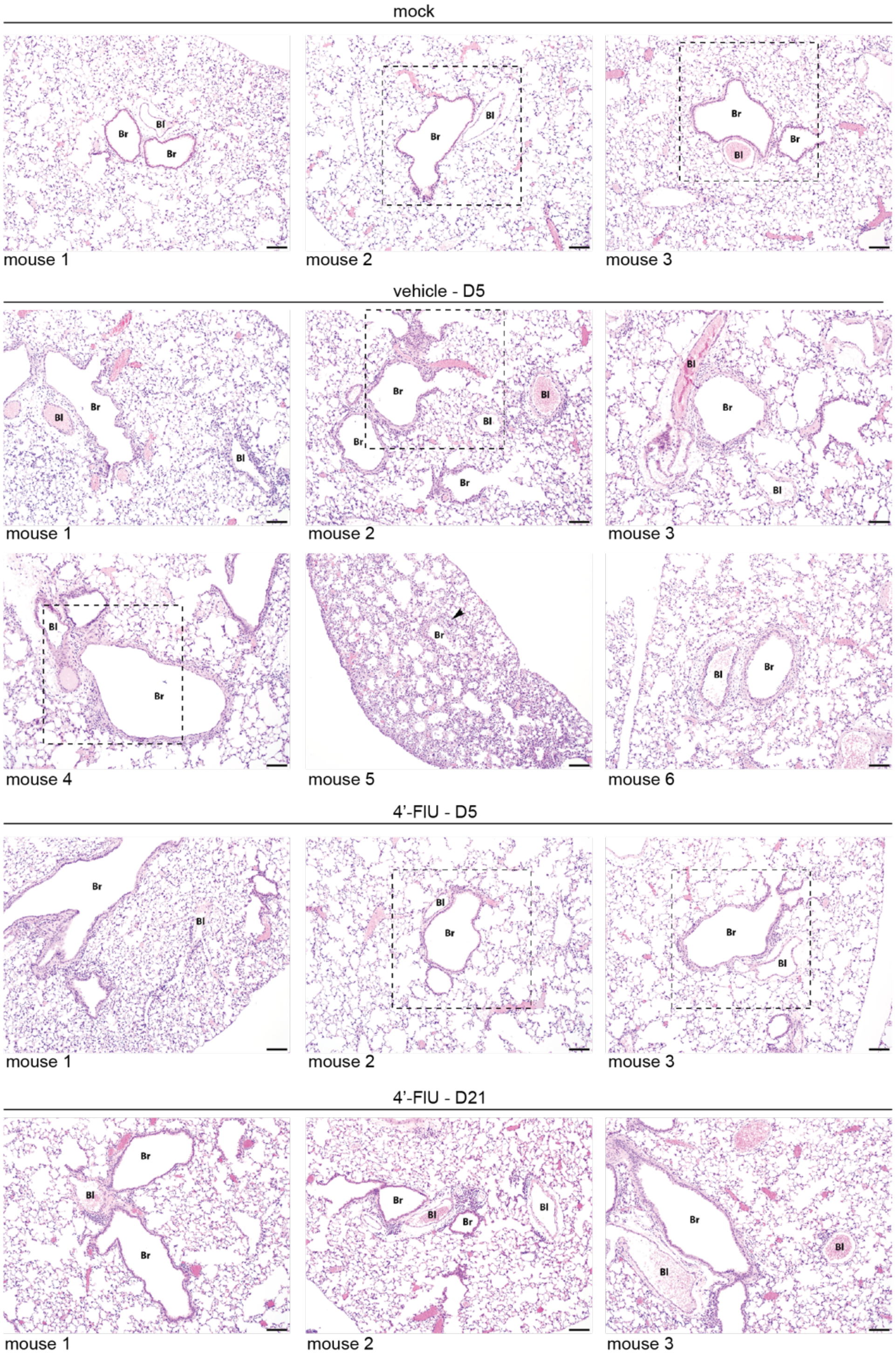


**S8 Figure:** Photomicrographs of lung tissues. Lungs were extracted on study days 5 or 21 (infected and treated animals only) of all animals examined in this study; tissue section were H&E stained; dashed rectangle shows sections presented in figure 4e.; magnification 10×; scale bar 100 µm; Br, bronchiole; Bl or arrowhead, blood vessel.

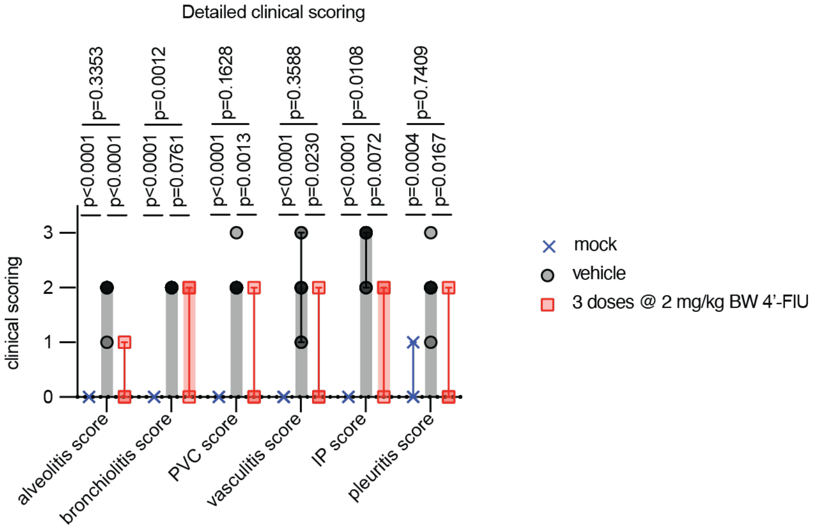


**S9 Figure:** Detailed histopathological scores of all animals examined in this study. Individual clinical scores for alveolitis, bronchiolitis, perivascular cuffing (PVC), vasculitis, interstitial pneumonia (IP), and pleuritis. Columns show data medians, symbols represent individual animals; 1-way ANOVA with Tukey's *post hoc* test; p values are specified; n values are specified in figure 4f.

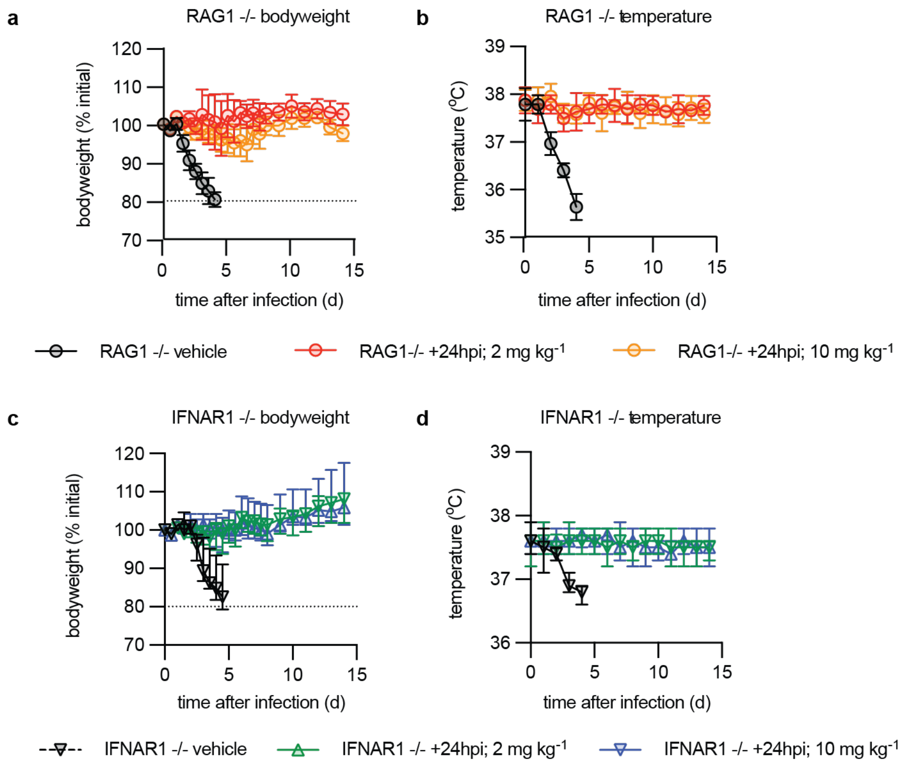


**S10 Figure:** Clinical signs in Ca09-infected immunocompromised mice treated with 4'-FIU. **a-c)** Shown are body weight measurements taken twice daily (a, c), and rectal body temperature (b, d) determined once daily. Symbols represent data means ± SD; n values are specified in figure 5. Results are shown for RAG1 KO (a-b ) and IFNAR1 KO (c-d) animals.

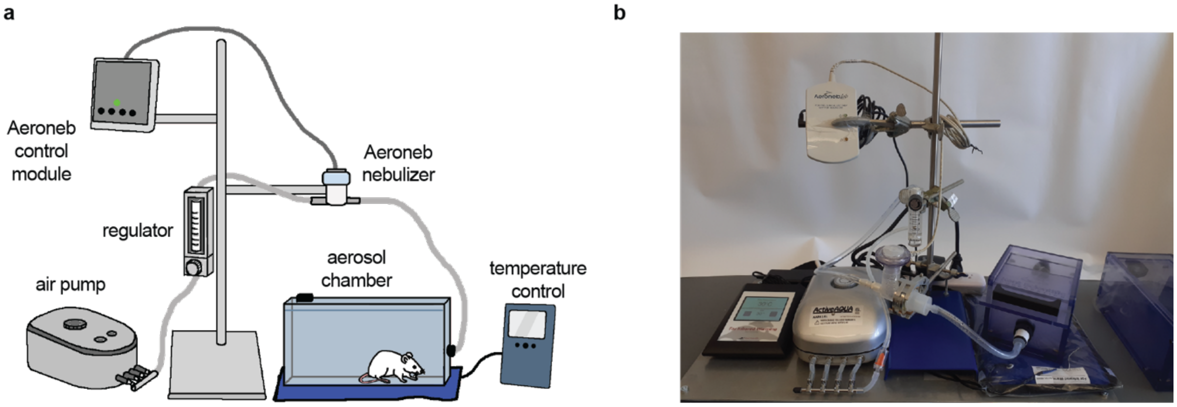


**S11 Figure:** Set-up of the Aeroneb nebulizer infection system. **a-b)** Schematic representation of the system (a) and photograph of the assembly (b).

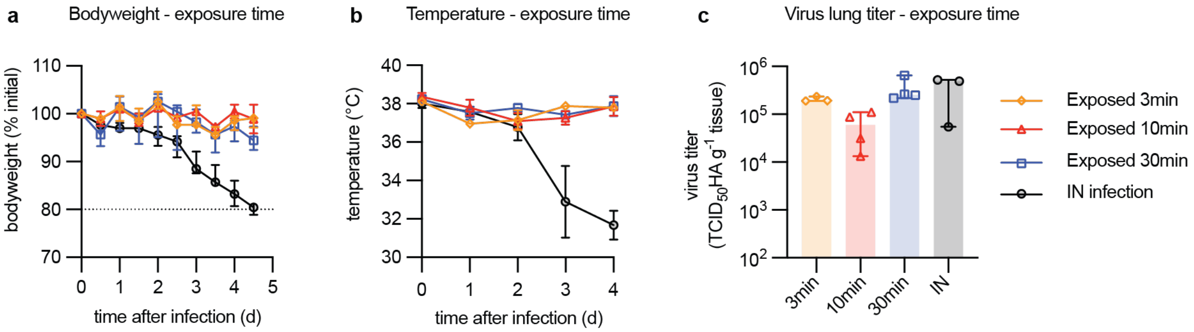


**S12 Figure:** S12. Variation of duration of animal exposure in the aerosolization chamber. **a-b)** Body weight measurements taken twice daily (a), and rectal body temperature (b) determined once daily. Symbols represent data means ± SD. **c)** Lung viral titers determined 4 days after infection. Columns represent data medians with 95% CI; symbols show individual animals; n=4.

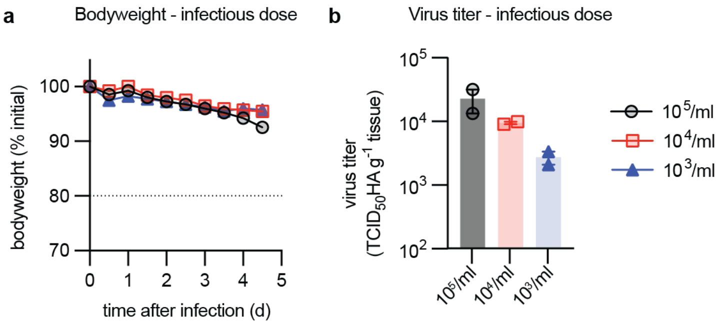


**S13 Figure:** Variation of Ca09 inoculum amount for infection in the aerosolization chamber. **a)** Body weight measurements taken twice daily. Symbols represent data means ± SD. **b)** Lung viral titers determined 4 days after infection. Columns represent data mean with range; symbols show individual animals; n=2.

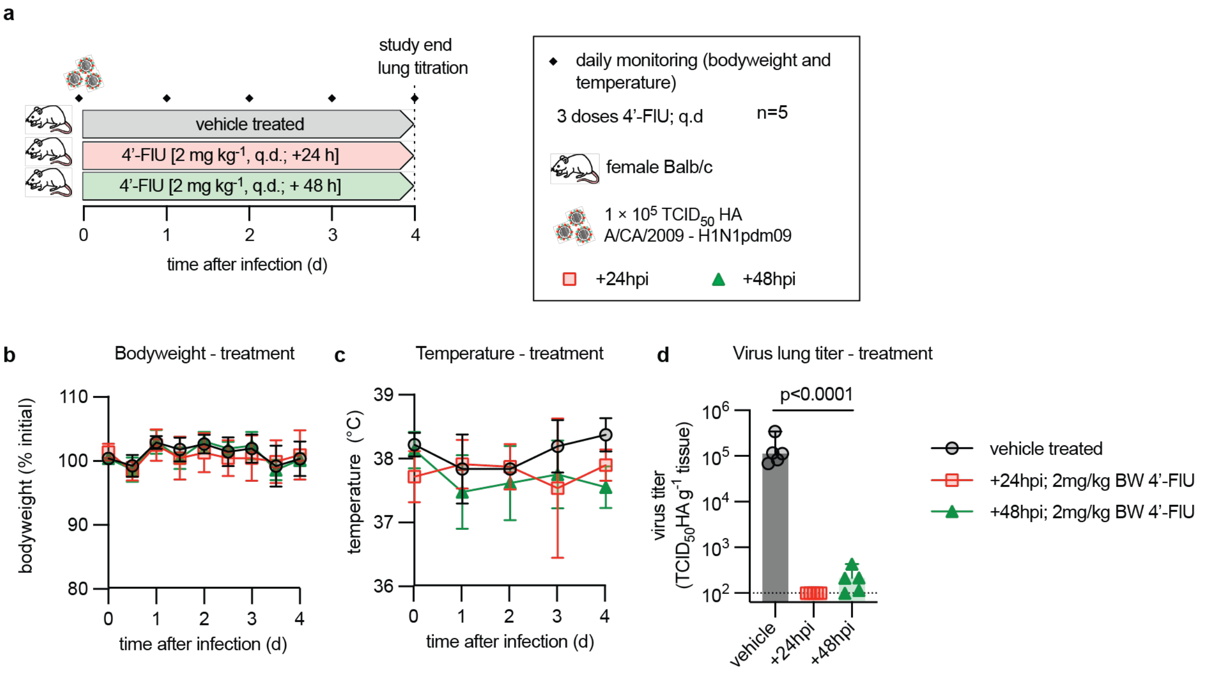


**S14 Figure:** 4'-FlU efficacy confirmation after infection with aerosolized inoculum virus. **a)** Efficacy study schematic; n values are specified. **b-c)** Body weight measurements of animals from (a) taken twice daily (a), and rectal body temperature (b) determined once daily. Symbols represent data means ± SD. **d)** Lung viral titers determined 4 days after infection. Columns represent data median with 95% CI, symbols show individual animals; 1-way ANOVA with Dunnett's *post hoc* test; P values are specified.

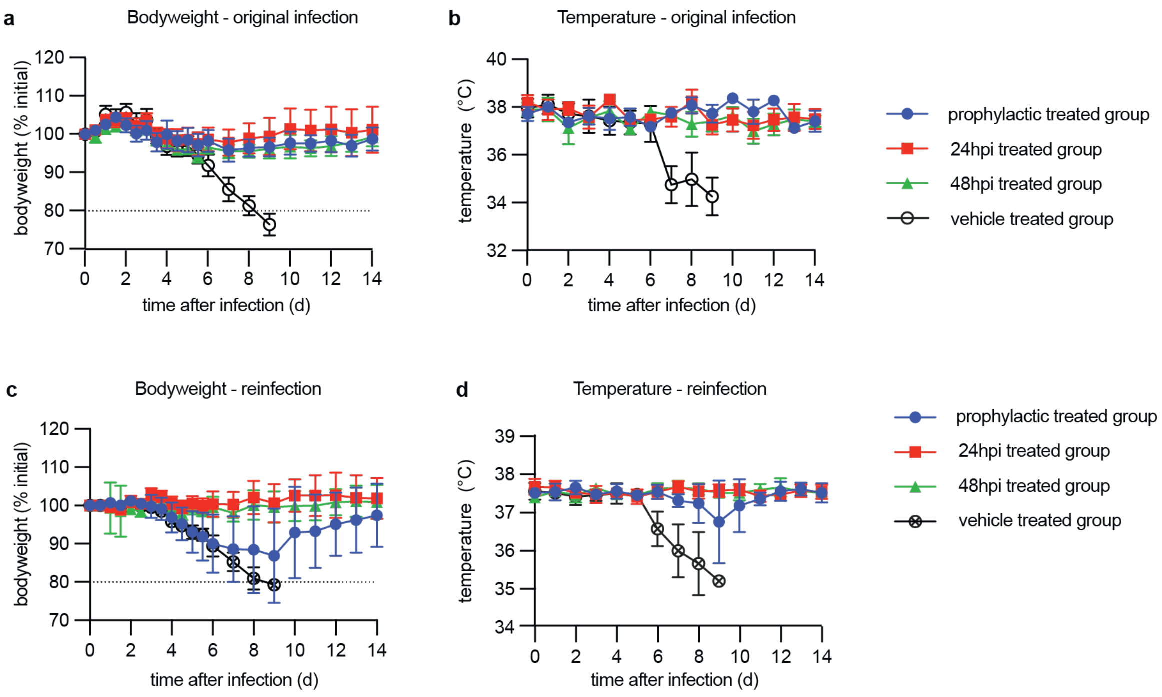


**S15 Figure:** Clinical signs of mice infected and reinfected with Ca09. **a-d)** Body weight measurements taken twice daily (a, c), and rectal body temperature determined once daily (b, d). Symbols represent data means ± SD. Results are shown for animals after the original infection (a-b) and after homotypic reinfection (c-d).

**S1 Data.** All quantitative source data generated in this study.

**S2 Data.** All statistical analyses performed for this study.
